# *Cryptococcus neoformans* capsule regrowth experiments reveal dynamics of enlargement and architecture

**DOI:** 10.1101/2021.11.15.468433

**Authors:** Maggie P. Wear, Ella Jacobs, Siqing Wang, Scott McConnell, Anthony Bowen, Camilla Strother, Radames J.B. Cordero, Arturo Casadevall

**Affiliations:** W. Harry Feinstone Department of Molecular Microbiology and Immunology, Johns Hopkins Bloomberg School of Public Health, Baltimore, MD, USA

**Keywords:** Sonication, Polysaccharide, Capsule, Cryptococcus, Glucanex

## Abstract

The polysaccharide capsule of fungal pathogen *Cryptococcus neoformans* is a critical virulence factor that has historically evaded characterization. Polysaccharides remain attached to the cell as capsular polysaccharide (CPS) or are shed into the surroundings in the form of exopolysaccharide (EPS). While a great deal of study has been done examining the properties of EPS, far less is known about CPS. In this work, we detail the development of new physical and enzymatic methods for the isolation of CPS which can be used to explore the architecture of the capsule and removed capsular material. Sonication and glucanex digestion yield soluble CPS preparations, while French Press and modified glucanex digestion plus vortexing remove the capsule and cell wall producing polysaccharide aggregates that we call ‘capsule ghosts.’ The existence of capsule ghosts implies an inherent organization that allows it to exist independent of the cell wall surface. As sonication and glucanex digestion were noncytotoxic, it was possible to observe the cryptococcal cells rebuilding their capsule, revealing new insights into capsule architecture and synthesis consistent with a model in which the capsule is assembled from smaller polymers, which are then assemble into larger ones.

**Importance:** Characterization of the cryptococcal polysaccharide capsule relies on methods of isolation for its *in vitro* study. This study demonstrates that the capsule is susceptible to physical and enzymatic removal. The application of new methods yields insights into the anatomy, modular nature, and architecture of the capsule with both soluble CPS preparations and ‘capsule ghosts.’ Together these insights inform on a long-standing debate modeling capsular assembly wherein our data shows that the capsule is assembled by smaller polymers added distally rather than by proximal addition or by polymers spanning the entire capsule radius.

## Introduction

The survival of *Cryptococcus* spp. in nature requires the yeast to defend itself against environmental stresses and phagocytic predators. Factors that afford this protection are hypothesized to function as virulence factors in the mammalian host (1–4). One such factor is the polysaccharide capsule of *C. neoformans* which protects the microbe from environmental desiccation (5) and amoeba predation (3). The capsule is comprised of two at least polysaccharides that have immunomodulatory and immunosuppressive activity: glucuronoxylomannan (GXM) and galactoxylomannan (GalXM) as well as mannoproteins at low abundance (6), though both GalXM and mannoproteins are hypothesized to only be secreted, not maintained in the capsule (7). Capsule size also plays a role in virulence and immune evasion (8). Larger capsule sizes are associated with more severe clinical outcomes as well a reduction in phagocytosis (9, 10). Establishing the essential nature of the capsule in pathogenicity, acapsular mutants exhibit a striking loss of virulence. (11, 12).

Cryptococcal polysaccharides are either attached to the cell as capsular polysaccharide (CPS) or are shed into the surroundings in the form of exopolysaccharide (EPS). Both CPS and EPS contribute to the immunosuppressive activity ascribed to the capsule, yet the two are antigenically and chemically distinct (13–15). A great deal is known about the properties of EPS due to well-established isolation protocols and ease of isolation. Until recently, most studies of EPS relied on cetyl trimethylammonium bromide (CTAB) extraction (16). CTAB is a detergent that can persist in the polysaccharide sample, hampering purification (13). Additionally, CTAB processing requires dehydration of the polysaccharide molecules, potentially collapsing or otherwise altering polysaccharide structure irreversibly. More recent work shows that EPS isolation can also be achieved by filtration methods that jellify the polysaccharide molecules, largely maintaining their hydrated, native state (13, 17).

In comparison with EPS, there is a relative dearth of literature describing CPS structure and function. The capsule is a highly hydrated structure composed primarily of water (Maxon). A major challenge of CPS isolation is its association with the cell wall. The most utilized methods are gamma irradiation and dimethyl sulphoxide (DMSO) extraction. Both methods have limitations. DMSO, like CTAB, persists in the sample after isolation and both the mechanism of action for capsule removal and how this may alter CPS structure are unknown. Gamma irradiation, which is thought to break glycosidic and other bonds through the radiolysis of water (6, 18–20), could result in the indiscriminate breakage of bonds, including the formation of open ring structures in the isolated polysaccharides (21, 22).

To address the need for new methodologies for the isolation of CPS we have explored physical and enzymatic methods of disrupting the linkage between the cell and capsule. This work describes three new methods of capsule disruption that add to the arsenal of capsular probing tools: ultrasonication, French Press, and glucanex digestion. Ultrasonication is a popular method used to perturb cells for lysis (23, 24). Ultrasonic sound waves (above 16 kHz) cause microscopic bubbles to form in solution. Collapse cavitation occurs when the bubbles implode resulting in high pressures and temperatures that can cause mechanical damage to surrounding materials (25). Stable cavitation is the consequence of oscillations in bubble size, this results in microstreaming (rapid flow of medium around the bubble), which produces sheer forces strong enough to damage macromolecules (25, 26). The French pressure cell press disrupts the *C. neoformans* capsule with a piston that exerts hydraulic pressure on the cell culture solution, forcing it through a narrow valve opening (27, 28). The principle of French press extorts fluid dynamics but with the application of high-pressure forces added to disrupt cell walls. Pressures of 25,000 psi have been utilized to lyse cells; however, far less pressure is necessary for capsular removal. Glucanex, or lysing enzymes from *Trichoderma harzianum*, contains β-1,3-glucanase, cellulase, protease, chitinase, and the more recently discovered α-1,3-endo-glucanase (29). This enzyme cocktail was utilized for the production of *C. neoformans* protoplasts, thus removing the cell wall, which in this work is shown to also include the capsule (30). In this work we use these new methods to gain new insights into capsule architecture.

## Results

### Removal of capsule by glucanex, sonication and DMSO

We evaluated the outcome of treating *C. neoformans* cells with sonication and glucanex and compared them to those treated with DMSO. Sonication resulted in a statistically significant reduction in capsule size in three *C. neoformans* strains, including those of serotype A and D. (Figure 1a). The starting capsule and cell body size varied with the strain, yet sonication efficiently removed material from all strains (Figure 1b, Supplemental Figure 1), leaving a median capsule radius after treatment of approximately 2 μm (Figure 1a, 2a). Glucanex is usually used in the cryptococcal field to generate spheroplasts as a prelude to lysing cells. Here we note that in removing the cell wall it also removes the polysaccharide capsule. For each of the three methods we measured: (i) reduction in capsule radius (ii) cell survival (iii) freed polysaccharide as measured by capture ELISA and Phenol Sulfuric Acid (PSA) assays and (iv) size of polymers in isolated CPS. Cells were analyzed for capsule-size reduction using Quantitative Capsule Analysis (QCA), a computer program developed in our laboratory that scans microscopic images to provide capsule dimension information (31). All three capsule removal methods produced a significant reduction in capsule size (Figure 2a), but DMSO killed the cells while both sonication and glucanex digestion were significantly less toxic, reducing viability to ^~^90% and ^~^60%, respectively (Figure 2b).

**Figure 1:**
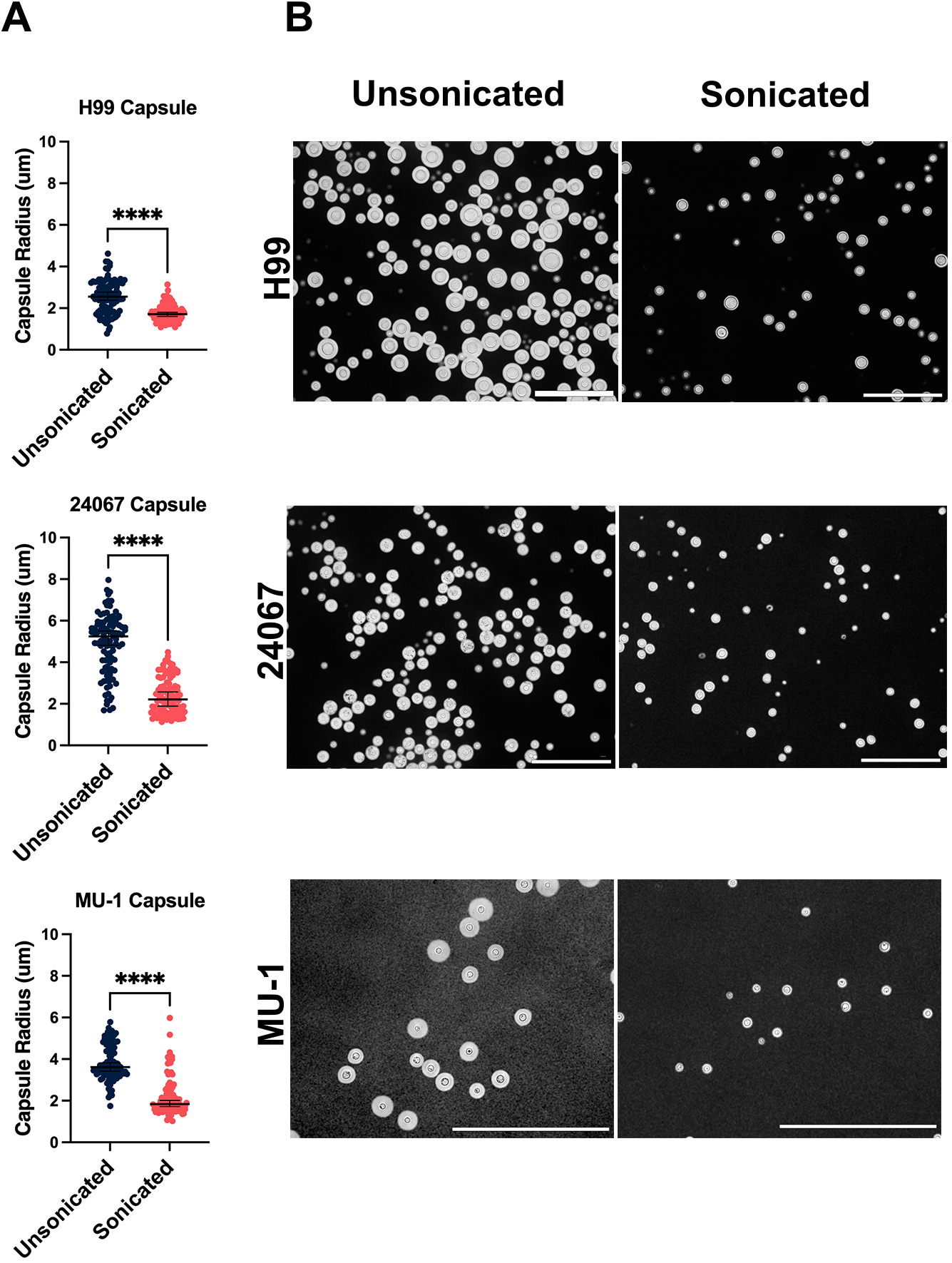
Effect of sonication on the capsule radius of *C. neoformans* serotype A and D strains. **A**. Quantitative capsule measurement before and after sonication in serotype A H99, serotype A single motif expressing strain Mu-1, and serotype D single motif expressing strain 24067. **B**. India ink microscopy images of strains before (left) and after (right) sonication. Scale bars represent 100 μm.

**Figure 2:**
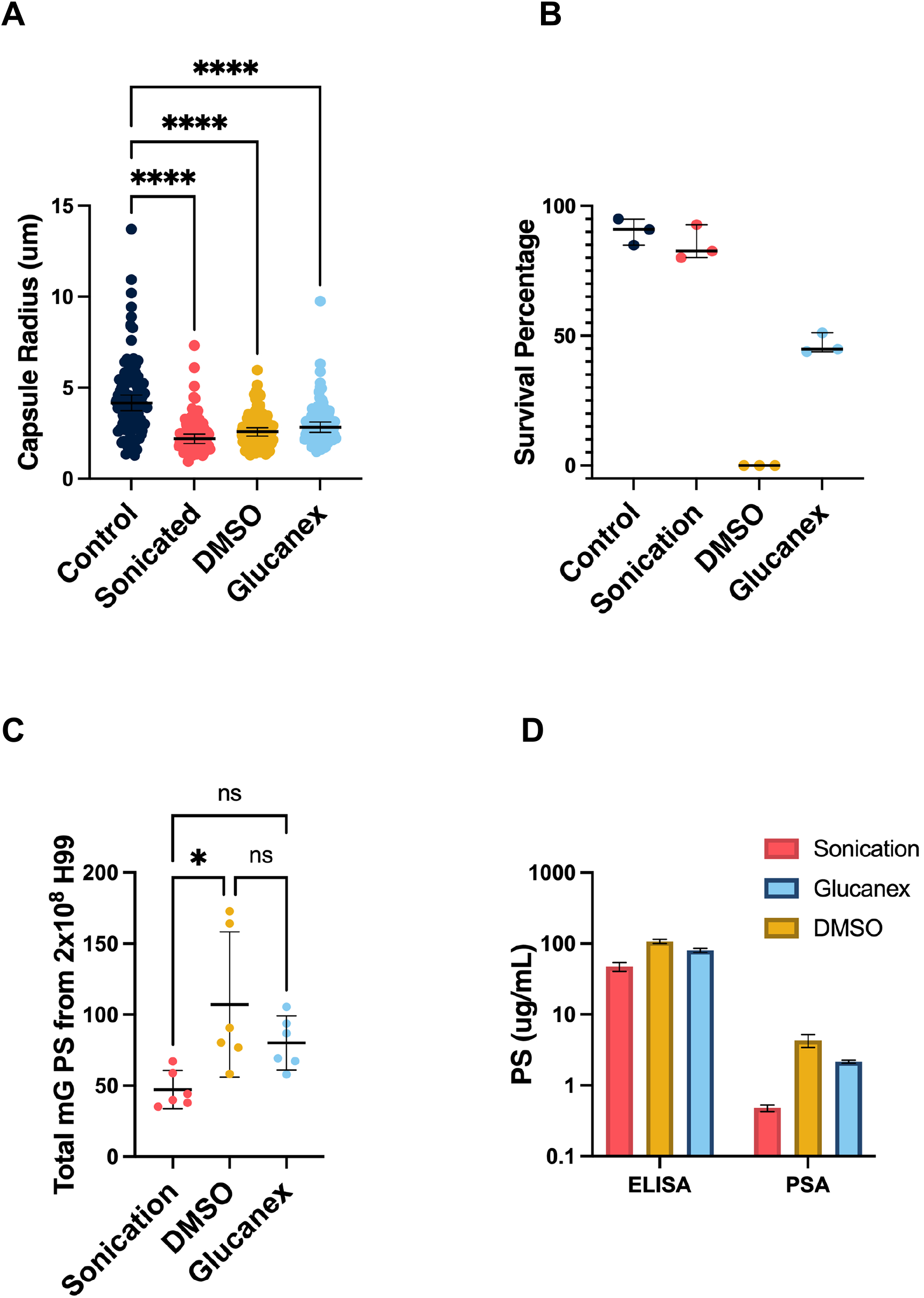
Comparison of capsular removal methods to impact radius, cell viability, and released polysaccharide. **A.** Quantitative measure of removal of capsule by sonication, DMSO, or glucanex treatment. **B.** Methylene blue survival of *C. neoformans* cells before (control) and after CPS removal. **C.** Total removed polysaccharide estimated by ELISA from 2×10^8^ H99 cells. Values represent three biological replicates with two technical replicates each. **D.** CPS yield as determined by capture ELISA and phenol sulfuric acid (PSA) assay, average of 3 biological replicates and standard deviation shown.

ELISA assays estimated a total polysaccharide yield from 2 × 10^8^ cells of 99 mg for DMSO, 78 mg for glucanex treatment, and 46 mg for sonication (Figure 2c). These polysaccharide yields are consistent with analysis of wet cell pellets before and after sonication, with the same number of cells yielding a total mass of 138 mg, which decreased to 98 mg after sonication, indicating that 44 mg of polysaccharide was released, a value that is close to that measured by ELISA. While capsule diameter is relatively negatively correlated with ELISA-measured supernatant GXM (r = −0.800, p = 0.333), PSA measurements of the same supernatant show less correlation with capsule diameter (r = −0.400, p = 0.7500). Thus, capsule diameter measurements and GXM quantification by ELISA are correlated, but PSA measurements are uncoupled. Since antibody-based detection methods are used in clinical and research settings, which rely on immunocomplexes between the GXM antigen and specific antibodies for detection and quantification, we utilized the capture ELISA data for quantification. Measurement of isolated CPS particle size showed high variance between biological replicates, but average effective diameter indicates that Glucanex digestion yielded the largest polymers while DMSO yields the smallest (Supplemental Figure 2). Additionally, CPS preparations by sonication or Glucanex treatment yielded a concentrated sample of CPS from 2 × 10^8^ cells in just 2 mL while the same number of cells results in 120 mL of preparation by DMSO.

### Glucanex and French press treatment produce capsule ghosts

During the processing and analysis described above, we observed what appeared to be capsule structures that no longer contained a cell. In an analogy to a technique to produce melanin ghosts (32), we refer to these structures freed of their cells as capsule ghosts (Figure 3). Capsule ghosts are produced by glucanex digestion, 500 psi in a French Press, or lateral shearing. All three methods result in abnormally shaped capsules, but French Press results in cells that are cracked open while Glucanex digestion and lateral shearing - produced by finger-pressing the encapsulated *C. neoformans* cells between a microscope slide and cover slip – results in capsular blebbing and capsules absent a cell but retaining the cell wall (as confirmed by staining with Uvitex2B) (Figure 3a). Further immunofluorescence analysis of the Glucanex-derived capsule ghosts showed that they did not contain cellular material, as revealed by the absence of DAPI staining, but did contain GXM as evident by reactivity with mAb 18B7 (Supp. Figure 3). The Glucanex-derived capsule ghosts also had polysaccharide reducing ends, as inferred from reactivity of aldehyde groups with a reducing end probe (hydroxylamine-488) (33), in the area where the capsule would have been in contact with the cell wall (Figure 3b). During optimization it was noted that yeast cell concentration was important for French Press. Cellular breakage easily occurred at a concentration of 1 × 10^7^ and a low pressure of 500 psi. We observed that cells at a concentration of 1× 10^8^ did not have a capsule size reduction when exposed to the French Press at 500 psi for two passes.

**Figure 3:**
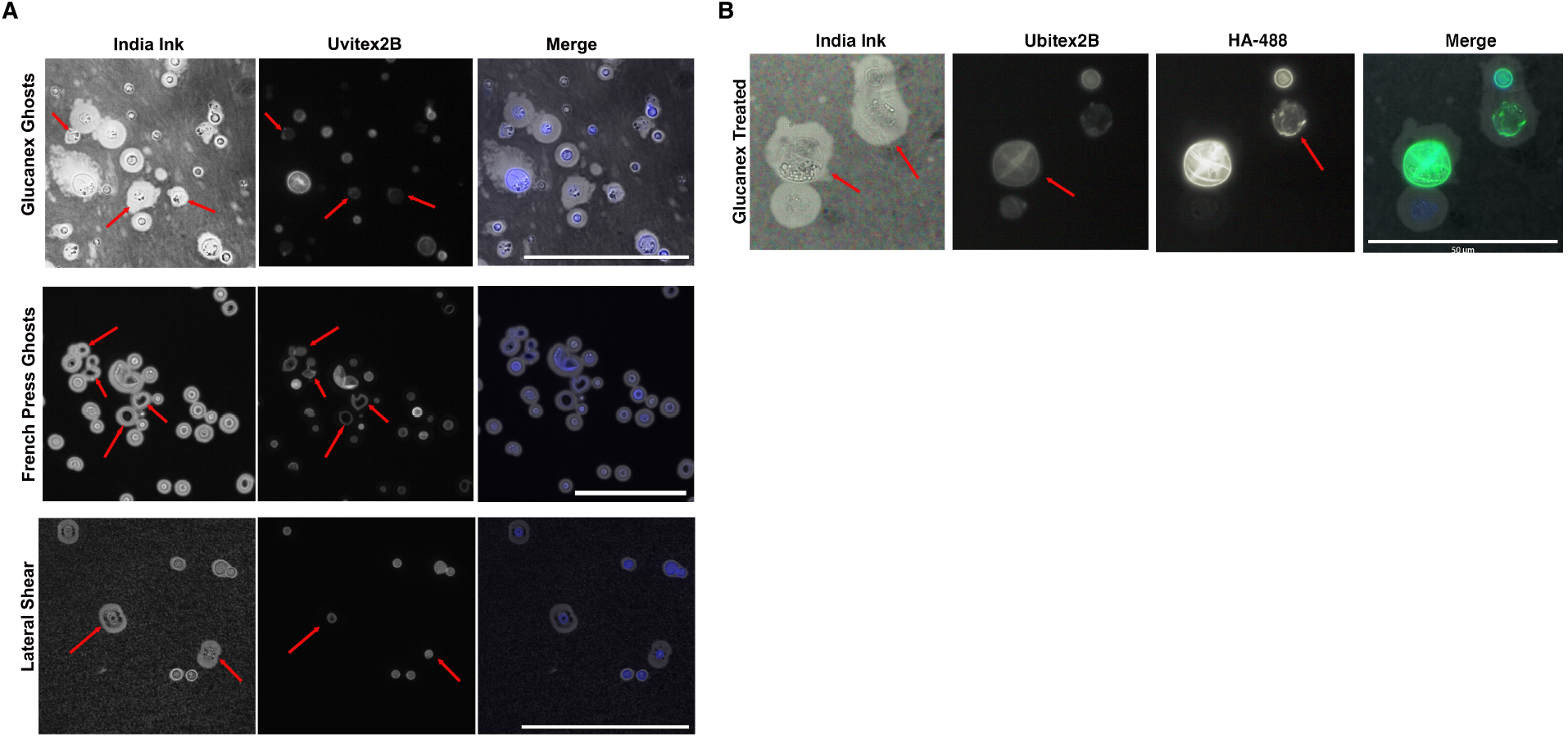
Glucanex, French press, and Lateral Shearing treatment produce capsule ghosts. **A**. India ink microscopy of capsule ghosts resulting from glucanex digestion (top), French Press (middle), or lateral shearing (bottom) show that capsule ghosts contain cell wall (as stained by Uvitex 2b) but are abnormal in shape, cracked open, blebbing, with off-center cells. Scale bars represent 100 μm **B**. India ink microscopy with a polysaccharide reducing-end probe (HA-488) and cell wall staining (Uvitex-2b) showing that reducing-ends are found at the cell wall even after the cell has been lost from ghosts. Scale bars represent 50 μm. Red arrows indicate capsule ghosts.

### Sonication treated cells regrow their capsules with reducing ends throughout the capsule

Noting that sonication and glucanex digestion removed capsules without killing *C. neoformans* provided the first opportunity for studying capsule regrowth. Prior studies of capsule growth were limited to measuring capsule enlargement (34–36) after placing cells in capsule enlarging conditions, while sonication and glucanex digestion provides the opportunity to study the repair of these structures after removal of large portions of polysaccharide. We observed that after 17 h the sonicated cells had completely regrown their capsules (Figure 4a) but glucanex treated cells took longer, requiring more than 74 h to regrow their capsules and not reaching the capsule size of the untreated cells (Figure 4b). We also evaluated where reducing end-containing polysaccharides were incorporated into the capsule after sonication by adding the HA-488 reducing end probe to the culture during capsule regrowth. After 18 h both the sonicated cells alone and with the reducing end had regrown their capsules, thought those with the HA-488 lagged behind the control cells (Figure 4a). Additionally, the reducing end probe staining was observed throughout the regrown capsules (Figure 4b, which is different from the lack of reactivity with mature capsules (33). The H99 cells were exposed to a second round of sonication and were able to regrow their capsules (Figure 4a).

**Figure 4:**
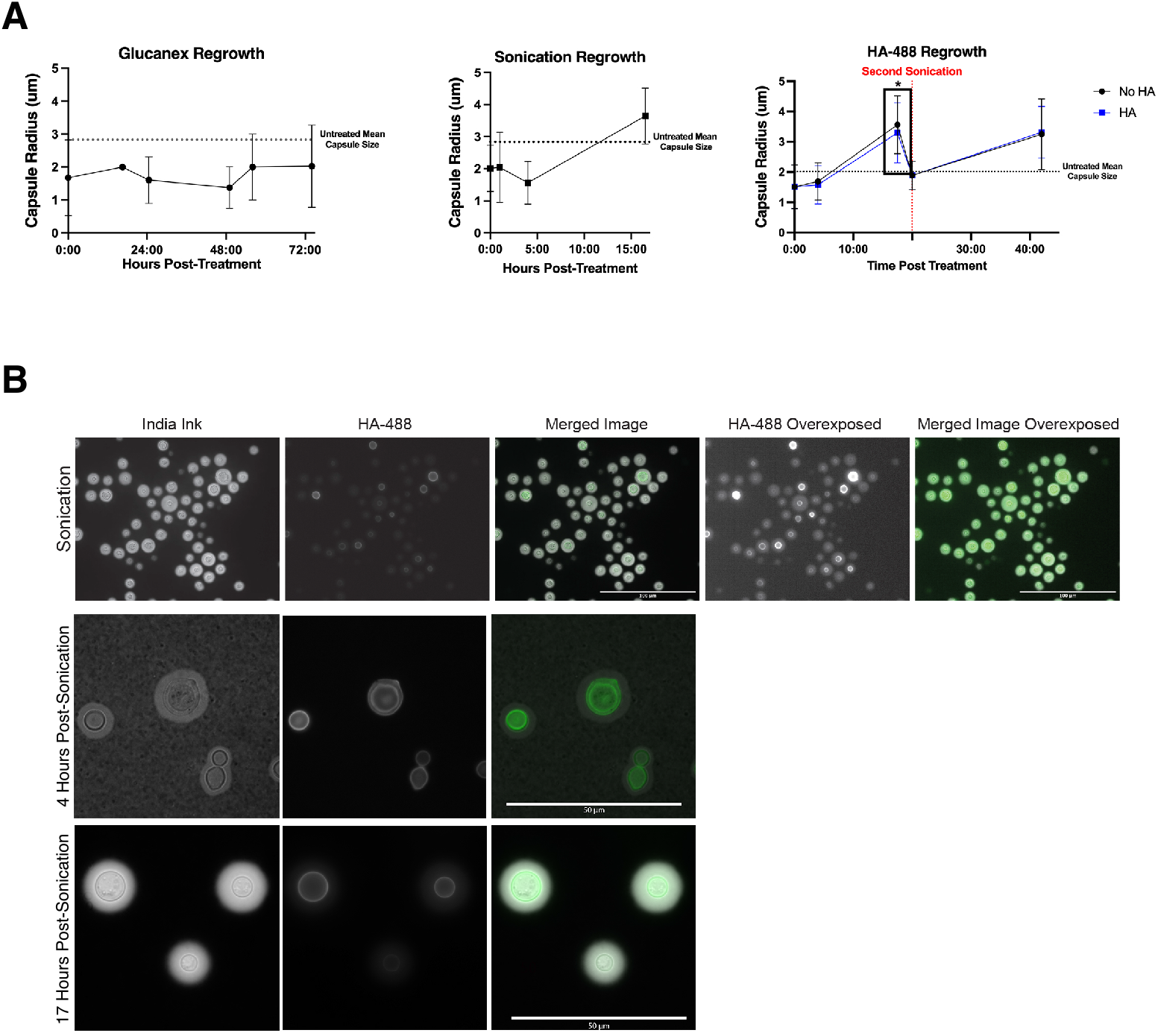
Examination of Capsule Regrowth after sonication and Glucanex digestion. For all experiments, before (untreated mean) after (0m) and at time intervals after cells were returned to minimal medial culture a 1mL aliquot was removed, washed, and imaged with India ink. The images from India ink staining were evaluated by QCA to determine median capsule size (listed on plot). **A.** Capsule growth after 76 hours post glucanex digestion (middle). Capsule growth 17 hours post sonication (middle). Capsule growth two sonication events in the presence or absence of HA-488 reducing end probe (right). The HA-488 stained cells show statistically significantly (unpaired student’s t-test, p-value <0.05) smaller capsules at 17 hrs. **B.** Immunofluorescence of India ink and reducing end staining after sonication. Overexposed panel shows that all cells stain with HA-488, but with different intensities. HA-488 staining is most intense at the cell wall-capsule interface, but is found throughout the capsule, even at 4 hours post sonication.

### Effects of sonication on antibody and complement binding and phagocytosis

To assess how the sonication-mediated removal of CPS affected the capsule of *C. neoformans* we performed immunofluorescence (IF) staining and microscopy with two different mAbs to GXM. While capsule reduction analysis shows that not all the capsule is removed by sonication, IF shows that the staining of mAbs 2D10 (IgM) and 2H1 (IgG) staining was less intense (Figure 5a). Both antibodies stained the capsule before and after sonication (Figure 5b), but after sonication there was a statistically significant loss of staining intensity as defined by integrated density. Staining for C3 was also statistically significantly reduced after sonication-mediated removal of CPS.

**Figure 5:**
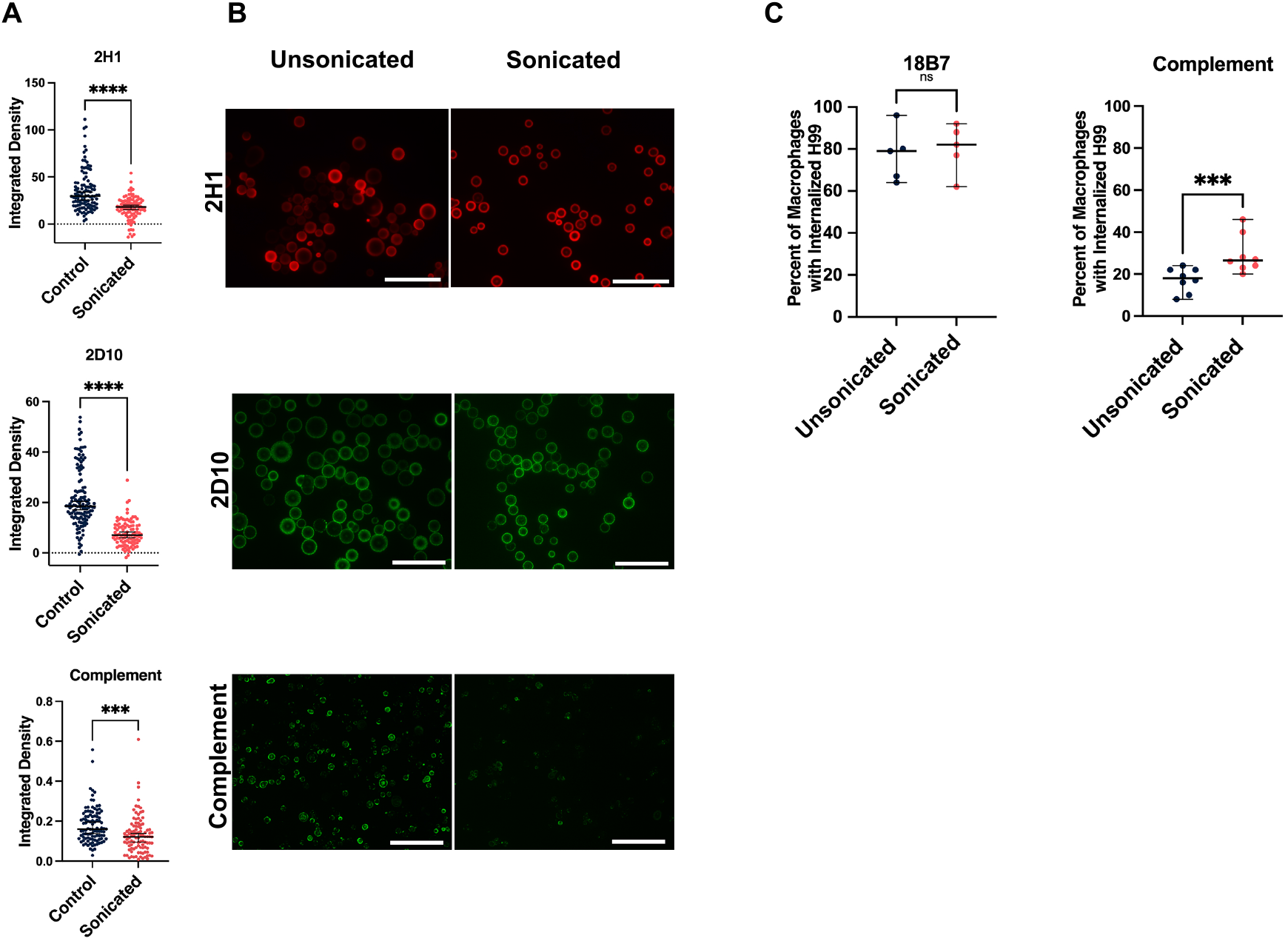
Effect of sonication on antibody and complement binding to the capsule. **A**. integrated density analysis of immunofluorescence staining of *C. neoformans* cells before and after sonication. **B**. Immunofluorescence microscopy images of strains before (left) and after (right) sonication with anti-GXM IgG 2H1 or IgM 2D10 or complement C3. Scale bars represent 100 μm. **C**. Percentage of BMDM cells with internalized cryptococcus via complement-mediated (left) or antibody mediated (right) mechanisms before and after sonication. Scale bars represent 100 μm.

When the sonicated cells were added to macrophages and the phagocytic index was measured, sonicated cells showed a statistically significant increase in phagocytosis in the presence of complement (Figure 4c). In contrast, no change in phagocytic efficacy was observed with antibody-mediated opsonization with mAb 18B7.

### Models of Capsule Assembly

Three major theories of capsular assembly have been proposed. Theory 1 asserts that material is added at the inner portion of the capsule, thereby dislocating existing polymers and moving them outwards from the cell (Figure 6a). This is supported by data with fluorescently labeled antibodies (37) and live imaging with antibodies (38). Theory 2 puts forth that capsule assembly occurs by incorporation of new material at the distal edge of the capsule (Figure 6b). This is supported by capsule regrowth after gamma irradiation (34). Theory 3 combines elements of Theories 1 and 2. In addition, a variation of theory 3 posits that the capsule is made of polymers which span the entire distance from the cell wall to the capsule edge (Figure 6c), at least for capsular enlargement. This is supported by the observation that polymer size and capsule radius are linearly correlated with polystyrene bead penetration (36) as well as recent experiments with the reducing end probe (33). Our observations showing reducing ends maintained at the cell wall-capsule interface in both the growing and mature capsule is consistent will all three theories. However, we also observe reducing ends throughout the capsule, but only when the reducing end probe is present during capsule assembly, suggesting that these are present during early assembly but blocked or not accessible in more mature capsules, a result that fits best with Theory 2. Finally, the size of polymers in CPS preparations do not show a linear correlation with the radius of capsule removed using these three methods, with the caveat that in this project we studied de novo capsule growth and not enlargement, as per the prior study, which used DMSO to remove CPS (36).

**Figure 6:**
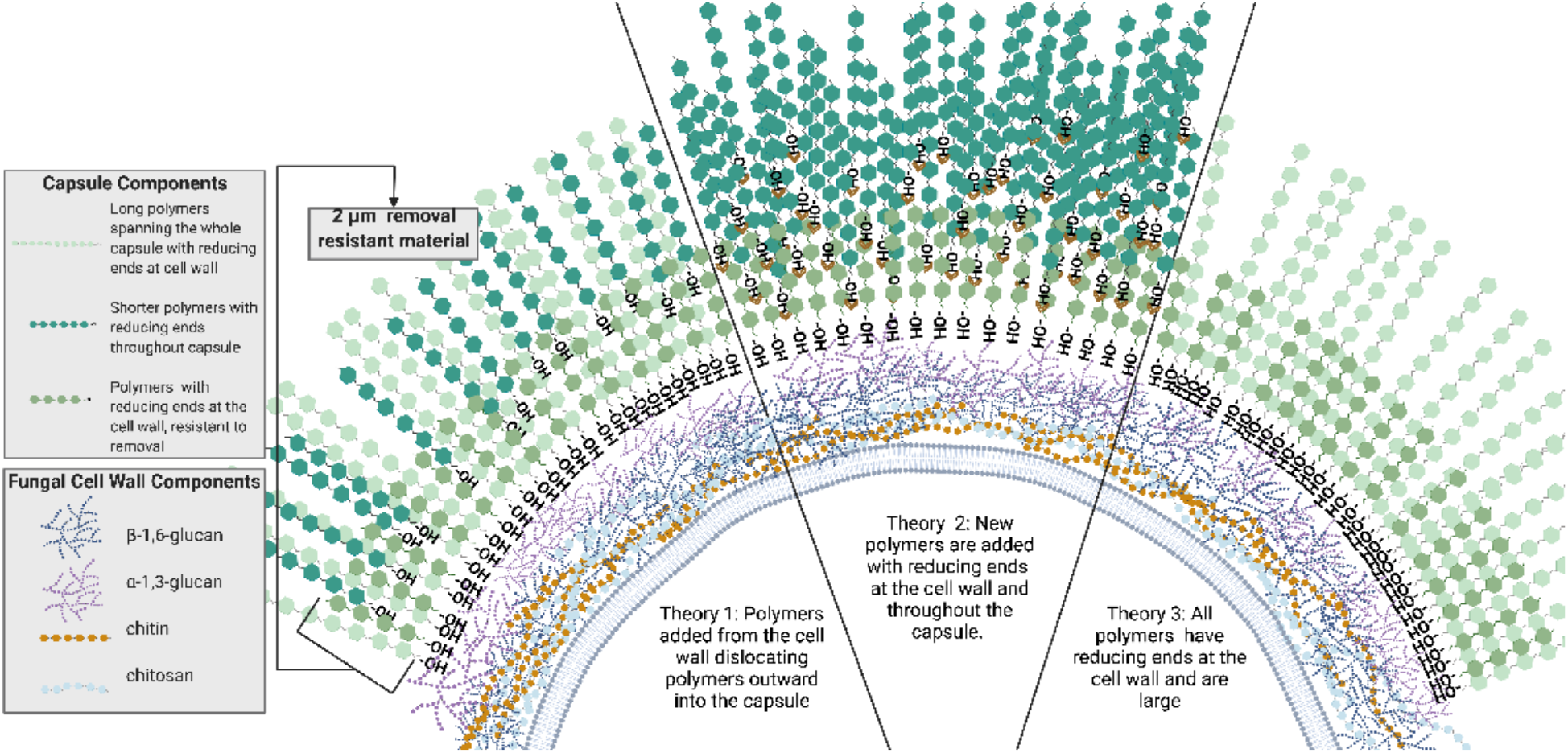
Models of capsule rebuilding. A diagram of three potential versions of capsule construction as informed by the data: 1: Polymers are added at the cell wall dislocated outward 2: New polymers are added both at the cell wall and throughout the capsule 3: All polymers remain at the cell wall and some transverse the entire capsule. 2 μm of removal-resistant material is present. Figure created with BioRender (54)

## Discussion

The capsule of *C. neoformans*, while critically important to fungal virulence, has eluded in depth analysis due to both its hydrated nature (39) and its attachment to the cell wall. Structural insights have generally come from extrapolating the data derived from EPS, but work in 2008 concluded that EPS and CPS are distinct from one another physically, chemically, and antigenically (13). We explored two physical methods – sonication and French press – and an enzymatic method, glucanex digestion, and compared these to chemical removal with DMSO. DMSO results in the highest yield of CPS (99 mg per 2×10^8^ cells), however, while both glucanex and sonication have lower yields (78 mg and 46mg respectively), the ease and quick turnaround time of these methodologies present a distinct advantage. Further, as a physical method that does not introduce a chemical or enzyme to the sample, sonication is the CPS isolation technique least likely to alter the native structure of the polymers.

Three methods, French press, modified glucanex digestion, and lateral shearing, resulted in the decapsulation of *C. neoformans*, but not into soluble polymers. India ink staining microscopy revealed that H99 cells exposed to relatively low pressures in the French Press (^~^500 psi) produced cells broken in half, but not fully stripped of capsular material. The cell wall was still present in these French-Press shells, as evidenced by Uvitex-2b staining. After treatment with glucanex and vortexing, entire capsules without a cell body were observed, and we are calling these structures capsule ghosts. This phenomenon may be attributable to the distinct cell wall architecture of cryptococcal cells. *C. neoformans* has less β-1, 3 glucan, a target of glucanex, than other yeasts and it is localized to the exterior layer of the cell wall (40). Additionally, in the conditions used to produce glucanex, *Trichoderma harzianum* also produces α-1,3-endo-glucanase which can directly cleave the α-1,3-glucan involved in GXM attachment to the cell wall (29, 41, 42). Repeated or excess pressure through lateral sheering can disrupt or break the cell body apart, resulting in off-center cell bodies, permeabilized capsules, and half cells like those observed during French Press treatment. The shearing process appears to impact capsule-cell wall interactions as similar phenomena have been observed in cells with partial inhibition of chitin synthesis (43)(Figure 5). Together these data suggest that the removal of the capsule can occur in two ways, one which produces soluble polysaccharide particles, and another which allows capsular architecture to remain intact after separation from the cell body. The fact that the capsule can be separated from the cell body means that it is held together in ways that are not dependent on its contacts with the cell wall, suggesting the existence of a network of intermolecular linkages that hold the polysaccharide molecules together.

Of the three methods for capsule removal, sonication was associated with the highest cell survival and was therefore used for further immunogenic characterization. We observed that both antibody and complement bound to the partially stripped capsules, albeit at reduced amounts. This is consistent with previous work showing that the innermost region of the capsule has the lowest antibody binding capacity while outer regions, removed by sonication, have higher predicted binding capabilities (6, 44). The patterns of antibody binding were unchanged by sonication, with the binding of mAb 2D10 remaining punctate before and after sonication, while mAb 2H1 exhibited sustained annular staining, consistent with previous findings (45). While the binding of mAbs 2H1 and 2D10 was decreased, the efficiency of antibody-mediated phagocytosis was not impacted by sonication. Even though complement deposition was decreased after sonication, sonicated H99 cells opsonized with complement had a greater percentage of phagocytic positive macrophages. Complement-mediated phagocytosis, unlike antibody-mediated phagocytosis (46), was previously shown to be inversely correlated with yeast cell size and is hypothesized to be impacted by deposition patterns (47). Together these data indicate that the inner ^~^2μm of the capsule not removed by sonication differs from the outer region in antibody binding and complement-mediated phagocytosis.

In addition to producing isolated CPS, each of these methods yielded new insights into capsular architecture. Based on the amount of CPS released we can estimate the density of the capsule removed by each method. Previous work using gamma irradiation-mediated CPS release showed capsule density lessens towards the periphery (35). This work revealed densities of ^~^110 μg/μl at a distance of ^~^1 μm from the cell wall which then decreased to 20 μg/μl at 1.5 μm, 10 μg/μl at 2.3 μm, and <5 μg/μl at 3 μm (35). Using the same method of quantification, our isolated CPS densities are 83 μg/μl for DMSO, 8 μg/μl for sonication, and 42 μg/μl for glucanex digestion (average values for the entire region spanning the outermost layers of the capsule to ~2 um from the cell wall). While sonication yields a density in line with that estimated by Maxson *et. al*., both DMSO and glucanex digestion result in densities higher than those calculated based on CPS removal with gamma irradiation. Further study of the secondary, tertiary, and even quaternary structure of the capsule will be necessary to determine the complex interplay responsible for the different observed densities of the capsule.

Previous work to characterize capsular geography has involved the use of mAbs to GXM to assess capsule growth (46) and to determine how new polysaccharide polymers are added to the capsule (47). However, the binding of these antibodies can affect capsule structure through crosslinking effects (45) and antibody binding location could vary if epitope changes position during capsule growth. Several different theories on capsule growth have been proposed. Some involve the intermingling of old and new polymers (48) while others suggest polymers extend from the cell wall to span the entire capsule (49). Our results here add to, and confirm, a model of capsule assembly where smaller polymers are assembled distal to the ^~^2μm inner capsule layer to create the capsule. The median induced capsule diameter is 6 μm before CPS removal and ^~^2 μm remains after the CPS has been isolated, resulting in ^~^4 μm of capsule removed. The effective diameter of isolated CPS as measured by DLS varied by preparation method – DMSO yields 0.28 μm, Sonication yields 0.4 μm, glucanex yields 0.7 μm, and glucanex ghosts yield 1.1 μm – but none of these were large enough to span the 4μm distance that was removed. When we examine the polysaccharide reducing ends using a fluorescent probe, they are only observed at the cell wall-capsule interface in capsule ghosts. However, when cells are decapsulated by sonication and the capsule is allowed to regrow in the presence of the reducing end probe we observe that the regrowth was slower. Additionally, the entire capsule exhibited dim fluorescence consistent with the presence of the probe, which in turns implies the presence of reducing ends in the young capsule. This observation contrasts with the finding that mature capsules have no reactivity with reducing end probes consistent with the notion that these are blocked or not accessible. Lactonohydrolases have been reported to modify the capsular polysaccharide in the exterior space to create more crosslinked and branched molecules (48). Collectively, this data suggests that CPS polymers removed by these methods were not large enough to span the entire capsule, that the addition of the reducing end probe retards capsule regrowth, and that there are free reducing ends during the early stages of capsule assembly.

Taken together the employment of these new CPS isolation methods provide new insights into capsular assembly. After treatment by sonication or glucanex digestion ^~^2 μm of capsule remains. This inner portion is denser, moderates complement-mediated phagocytosis, and is the anchor point for reducing ends of some capsular polysaccharides. Reviewing previous studies, we note that the inner ^~^2 μm of capsule was also found after gamma-irradiation (44) as well as in nutrient-rich conditions (47), implying the existence of a more tenacious inner layer that appears to essential for capsule assembly. Together these data suggest that like the regulation of the maximum size of the capsule (49), there may also be a retained size for the capsule of ^~^2 μm, at least for the H99 strain. Further, the observation that there is no correlation between *in vitro* capsule size and virulence (50–52) is supported by our observation that the capsule can be regrown, suggesting that rather than size, the capsule’s critical virulence factor is its presence or absence, and the specific GXM polymer that is expressed. Additionally, our results are most consistent with the theory of capsule architecture where polymers are exported from the cell with free reducing ends and during incorporation into the capsule these reducing ends are made unavailable for binding, possibly through crosslinking as previously proposed (53). In this model polysaccharides are smaller in size and linked together or self-aggregate to span the entire capsule radius.

In summary, the application of physical, chemical, and enzymatic methods of decapsulation yield methods for both soluble CPS and in-tact “capsule ghosts” that minimally perturb the native capsular architecture. In addition, while sonication decreases complement binding, complement-mediated phagocytosis is increased in sonicated cells which could indicate that the remaining inner 2μm of capsule is more immunogenic than the outer region that was removed. Finally, we observed that decapsulated cells can regrow their capsule and this young capsule contains free polysaccharide reducing ends throughout suggesting capsule architecture evolves as the capsule ages.

## Experimental procedures

### Yeast strains and Culture

In capsule measurement, immunofluorescence, French Press, and phagocytosis experiments, H99 (serotype A), single expressing motif strain MU-1 (serotype A), and 24067 (serotype D) cells were sub-cultured into 6 mL Sabouraud liquid media for two days at 30°C on a culture wheel. Cells were washed twice in PBS and 3 mL of the culture was sub-cultured into 500 mL minimal media in a tissue culture flask at 30°C with shaking (150 RPM). (15 mM D-glucose, 10 mM MgSO4• 7 H2O, 20.3 mM KH2PO4, 3 mM Glycine, 10 mg/mL Thiamine pH 5.5). For capsular material removal experiments H99 (serotype A) was cultured in 5 mL YPD media for two days at 30°C on a culture wheel. The cells were subsequently washed twice with PBS, and 500 μl was sub-cultured into 500 mL minimal media in a tissue culture flask at 30°C with shaking (150 RPM). All cultures used grew in minimal media for 3-5 days. Strains were preserved in 15% Glycerol and maintained at −80°C.

### India Ink/QCA

*C. neoformans* cells were mixed with India Ink and imaged on an Olympus AX70 microscope using QImaging Retiga 1300 digital camera and the QCapture Suite V2.46 software (QImaging). Capsule measurements were produced using the exclusion zone produced with India Ink and the Quantitative Capture Analysis program developed by the lab (31). A minimum of 100 yeast cells were measured for each strain and condition.

### Sonication

Cells cultured in minimal media for 3-5 days were washed 3 times with PBS and brought to a density of 1×10^7^ cells/mL. For capsule removal experiments cells were brought to a density of 1×10^8^ cells/mL. After washing, 2 mL of cell suspension in PBS were sonicated on ice using a horn sonicator (Fisher Scientific Sonic Dismembrator F550 W/ultrasonic Convertor) on either setting 3 or 7 for 30 seconds (8 and 17 watts, respectively). Sonicated cells used in phagocytic index experiments were washed post-sonication to remove CPS.

### French Press

H99 cells cultured in minimal media for 3-5 days were washed 3 times with PBS and brought to a density of 1×10^7^ cells/mL. 10 mL of culture was run through the French Press G-M^®^ High Pressure Standard Cell twice at a pressure of (500 psi). After passing cells through the French Press they were stained with 0.05% Uvitex 2B and 1 μl of culture was combined with 7 μL India Ink and imaged with Olympus AX70 microscope using QImaging Retiga 1300 digital camera and the QCapture Suite V2.46 software (QImaging).

### Glucanex digestion

Glucanex enzyme (Lysing Enzymes from *Trichoderma harzianum*) was prepared at a concentration of 50 mg/mL in spheroblasting buffer (1M sorbitol, 10 mM EDTA, and 100 mM sodium citrate). H99 cells cultured in minimal media for 3-5 days were washed 3 times with PBS and brought to a density of 1×10^8^ cells/mL. 1 mL of cells were combined with 5 mL of glucanex enzyme and incubated for 2.5 hrs at 30°C while shaking. Cells were vortexed for 30 seconds on high or sonicated for 10 sec on power 3 and then spun 7 min at 1200 RPM to isolated capsule ghosts.

### DMSO treatment

500 μL of washed H99 cells at a density of 1×10^8^ cells/mL were added to 15 mL of pure Dimethyl sulfoxide (CH_3_)_2_SO. The cells and DMSO incubated for 30 minutes. At this time, the H99 cells were spun down and resuspended in 15 mL fresh DMSO. After 30 minutes the H99 cells were spun down a second time. 30 mL total DMSO with capsular material was recovered. The remaining DMSO was removed from the sample by dialysis with at least 10-12 solvent changes.

#### Capture ELISA

Microtiter plates were coated with anti-IgM Fc mAbs at a 1:1000 dilution in PBS at 50 ul per well. Plates were incubated at 37 degrees for 1 hour, then blocked with 200 ul blocking solution (1% BSA). After one hour of blocking at 37 degrees, plates were emptied by blotting over paper towels. Anti-GXM 2D10 IgG mAbs at 3 ug/ml, 50 ul per well were added. Plates were incubated at 4 degrees overnight and washed 3 times with TBS with 0.1% TWEEN-20 before use. An EPS standard was prepared at a top concentration of 10 ug/ml, then serial-diluted to make a total of 8 points. CPS supernatants obtained from DMSO, sonication, and glucanex treatment were diluted 500 times for the top concentration, then serial-diluted to make a total of 8 points. Serial dilutions were performed with blocking solutions. 50 ul of EPS standard and samples at different dilutions were added, and blocking solution was used as a negative control. Plates were incubated at 37 degrees for 1 hour and then washed 3 times. Anti-GXM 18B7 IgM mAbs at 5 ug/ml, 50 ul/well were added, plates were incubated for 1 hour and then washed 3 times. Alkaline phosphatase labeled anti-IgG Abs at 1 ug/ml, 50 ul/well were added, plates were incubated for 1 hour and washed 5 times. Finally, 1 mg/ml, 50 ul/well alkaline phosphatase substrate in substrate buffer were added. Plates were developed at RT and read at A405 until the top concentration standard absorbances reached just over 1. EPS A405 absorbances were plotted against concentrations and then fit with a linear model. This linear relationship was used to estimate the concentrations of CPS samples with respect to their A405 absorbances.

### PSA assay

This assay was developed from Maskuko et al., 2005. To start 100 μL of 1M mannose was 1:2 serially diluted into 50 μL of sterile water in the first three columns of a 96-well plate. Fifty microliters of each sample were put into three wells undiluted. A volume of 50 μL of sterile water was placed in three wells to determine background. One hundred microliters of concentrated sulfuric acid were placed in each well. Thirty microliters of a 5% phenol solution were pipetted into each well. The 96-well plate was then incubated at 37 °C for about 10 minutes. Plate was read at read at 490 nm.

### Statistical correlations of capsule degradation measurements

For each experimental capsule removal experiment (CPS removal or antibody-mediated capsule degradation), polysaccharide concentration in the supernatant was measured by capture ELISA, PSA and dry weight measurements. The total polysaccharide amount was then calculated to correct for differences in total volume. The corresponding CN cells were also analyzed after CPS removal or antibody treatment to measure the average capsule diameter. These measured values were then plotted as a scatter plot with respect to two variables at a time and the correlation coefficient by the Spearman nonparametric correlation test was calculated in GraphPad Prism 9.2.0.

### Immunofluorescence

To visualize the antibody staining, 5×10^6^ total cells were resuspended in 1mL blocking solution (prepared fresh, 1% BSA in 100 mL PBS) with 10 μg/mL of either of the primary mAbs that detect different epitopes within the PS capsule of *C. neoformans*. This study compared one IgM mAb, 2D10, and one IgG mAb, 2H1, antibodies. Complement binding changes were assessed by incubated cells in blocking buffer and 20% complement (guinea pig complement, MilliPore). H99 cells with either mAb or complement were placed on a shaker for 1 hr at 37°C. After primary opsonization, cells were spun down and washed with PBS then resuspended in 1mL blocking solution with either 1:500 Goat anti-Mouse IgM Heavy Chain Secondary Antibody conjugated to AlexaFluor 488 (Invitrogen), 1:500 Rabbit anti Mouse IgG (H+L) Secondary Antibody, Alexa Fluor 594, Invitrogen, or 1:50 goat anti-Complement C3 polyclonal—FITC (Invitrogen). H99 cells with either fluorescently conjugated anti-IgM mAb, anti-IgG mAb, or fluorescently conjugated anti-complement mAb were placed on a shaker for 1 hr at 37°C. Postincubation cells were washed once in PBS and imaged with Pro-Long Gold mounting solution (Molecular Probes). FITC channel excitation/emission was 498 and 516 nm respectively and TRITC excitation/emission was 540 and 580 nm respectively. Exposures of 2H1, 2D10, and complement binding were 100 ms, 400 ms, and 700 ms respectively. The same exposure was applied to both control and sonicated images. Images were collected with an Olympus AX70 microscope, photographed with a QImaging Retiga 1300 digital camera using the QCapture Suite V2.46 software (QImaging). Quantification of cells was achieved with ImageJ from Fiji (NIH) by measuring integrated density around each cell and subtracting background intensity from each measurement. No adjustment of brightness or contrast was performed for figure 3. Color filters were assigned with ImageJ from Fiji (NIH). All images in figure 5 were simultaneously brightened to the same degree in photoshop to view Uvitex2B/India ink contrast. Control and sonicated cells were measured with an area of the same size and the subtracted background was additionally scaled to this size.

### Phagocytic Index

Bone marrow-derived macrophages (BMDM) were obtained from 6-week-old C57BL/6 female mice. After 5-7 days of differentiation, the cells were seeded 5×10^4^ cells per Mat-Tek dish and allowed to settle for 30 minutes. After 30 minutes the dish was supplemented with 2 mL media. BMDMs were activated overnight with 0.5 μg/mL lipopolysaccharide (LPS, MilliporeSigma) and 10 ng/mL IFN-γ (Roche) for M1 polarization. BMDMs were maintained at 37 degrees C 9.5% CO_2_. *C. neoformans* were prepared and sonicated as described above. 1.5 × 10^5^ H99 were added for an MOI of 3. Opsonized cells were prepared with 10 μg/mL 18B7, or 20% complement (Fisher Scientific #642831) and placed on the slide portion of the dish. After a two-hour infection at 37 degrees C 9.5% CO_2_, BMDM cells were washed 4x with media to remove extracellular *C. neoformans* cells. Cultures were visualized on a Zeiss Axiovert 200M inverted microscope with a 10x phase objective in an enclosed chamber at 9.5 % CO2 and 37 °C. 100 BMDMs of each condition were counted and then scored for the presence of *C. neoformans* cells.

### Capsule Regrowth after Sonication and Glucanex Digestion

H99 cells were cultured, brought to a final concentration of 10^8^ cells/ml, and 2 ml cells were sonicated and treated with glucanex followed by sonication as described above. CPS supernatants were obtained after centrifugation, and treated cells were resuspended in 20ml minimal media and incubated with shaking at 30 ^o^C for capsule regrowth. India Ink slides were imaged and analyzed by QCA at several timepoints; 1 ml cells were aliquoted out and concentrated into 10 ul PBS for imaging. The sonicated cells were re-sonicated after approximately 48 hours of regrowth; half of the treated cells were directly resuspended in 20 ml minimal media for a second regrowth, and the other half were stained by a reducing end probe by adding 10ul HA-488 to 20ml minimal media into which the cells were resuspended. India Ink slides were imaged under white light and green fluorescence to track capsule regrowth and reducing end localization.

## Data Availability

All data are contained within the manuscript. Strains used in this study are available by request of the corresponding author.

## Supporting Information

This article contains supporting information.

## Acknowledgements

We would like to thank Conor Crawford for his work on the development of the reducing end probe for polysaccharides. The model figure 6 was created using BioRender. We would like to thank the MMI microscopy core for training and use of microscopy equipment for this work.

## Funding and additional information

This content is solely the responsibility of the authors and does not necessarily represent the official views of the National Institutes of Health. S.A.M. and M.P.W. were supported in part by NIH Grant AI007417. E.J. was supported in part by NIH grant AI138953-01AI.A.C. was supported in part by NIH Grants AI052733-16, AI152078-01, and HL059842-19.

## Bibliography

1. Casadevall, A. (2008) Evolution of intracellular pathogens. Annu. Rev. Microbiol. 62, 19–33

2. Casadevall, A., and Pirofski, L. (2003) The damage-response framework of microbial pathogenesis. Nat. Rev. Microbiol. 1, 17–24

3. Steenbergen, J. N., Shuman, H. A., and Casadevall, A. (2001) Cryptococcus neoformans interactions with amoebae suggest an explanation for its virulence and intracellular pathogenic strategy in macrophages. Proc. Natl. Acad. Sci. USA 98, 15245–15250

4. Colombo, A. C., Rella, A., Normile, T., Joffe, L. S., Tavares, P. M., de S Araújo, G. R., Frases, S., Orner, E. P., Farnoud, A. M., Fries, B. C., Sheridan, B., Nimrichter, L., Rodrigues, M. L., and Del Poeta, M. (2019) Cryptococcus neoformans Glucuronoxylomannan and Sterylglucoside Are Required for Host Protection in an Animal Vaccination Model. MBio 10

5. Aksenov, S. I., Babyeva, I. P., and Golubev, V. I. (1973) On the mechanism of adaptation of micro-organisms to conditions of extreme low humidity. Life Sci. Space Res. 11, 55–61

6. Zaragoza, O., Rodrigues, M. L., De Jesus, M., Frases, S., Dadachova, E., and Casadevall, A. (2009) in Advances in Applied Microbiology pp. 133–216, Elsevier

7. Vartivarian, S. E., Reyes, G. H., Jacobson, E. S., James, P. G., Cherniak, R., Mumaw, V. R., and Tingler, M. J. (1989) Localization of mannoprotein in Cryptococcus neoformans. J. Bacteriol. 171, 6850–6852

8. Vecchiarelli, A., Pericolini, E., Gabrielli, E., Kenno, S., Perito, S., Cenci, E., and Monari, C. (2013) Elucidating the immunological function of the Cryptococcus neoformans capsule. Future Microbiol 8, 1107–1116

9. Robertson, E. J., Najjuka, G., Rolfes, M. A., Akampurira, A., Jain, N., Anantharanjit, J., von Hohenberg, M., Tassieri, M., Carlsson, A., Meya, D. B., Harrison, T. S., Fries, B. C., Boulware, D. R., and Bicanic, T. (2014) Cryptococcus neoformans ex vivo capsule size is associated with intracranial pressure and host immune response in HIV-associated cryptococcal meningitis. J. Infect. Dis. 209, 74–82

10. Yasuoka, A., Kohno, S., Yamada, H., Kaku, M., and Koga, H. (1994) Influence of molecular sizes of Cryptococcus neoformans capsular polysaccharide on phagocytosis. Microbiol. Immunol. 38, 851–856

11. Chang, Y. C., and Kwon-Chung, K. J. (1994) Complementation of a capsule-deficient mutation of Cryptococcus neoformans restores its virulence. Mol. Cell. Biol. 14, 4912–4919

12. McClelland, E. E., Bernhardt, P., and Casadevall, A. (2006) Estimating the relative contributions of virulence factors for pathogenic microbes. Infect. Immun. 74, 1500–1504

13. Frases, S., Nimrichter, L., Viana, N. B., Nakouzi, A., and Casadevall, A. (2008) Cryptococcus neoformans capsular polysaccharide and exopolysaccharide fractions manifest physical, chemical, and antigenic differences. Eukaryotic Cell 7, 319–327

14. Kozel, T. R., Gulley, W. F., and Cazin, J. (1977) Immune response to Cryptococcus neoformans soluble polysaccharide: immunological unresponsiveness. Infect. Immun. 18, 701–707

15. Pericolini, E., Cenci, E., Monari, C., De Jesus, M., Bistoni, F., Casadevall, A., and Vecchiarelli, A. (2006) Cryptococcus neoformans capsular polysaccharide component galactoxylomannan induces apoptosis of human T-cells through activation of caspase-8. Cell Microbiol. 8, 267–275

16. Cherniak, R., Valafar, H., Morris, L. C., and Valafar, F. (1998) Cryptococcus neoformans chemotyping by quantitative analysis of 1H nuclear magnetic resonance spectra of glucuronoxylomannans with a computer-simulated artificial neural network. Clin Diagn Lab Immunol 5, 146–159

17. Nimrichter, L., Frases, S., Cinelli, L. P., Viana, N. B., Nakouzi, A., Travassos, L. R., Casadevall, A., and Rodrigues, M. L. (2007) Self-aggregation of Cryptococcus neoformans capsular glucuronoxylomannan is dependent on divalent cations. Eukaryotic Cell 6, 1400–1410

18. Bryan, R. A., Zaragoza, O., Zhang, T., Ortiz, G., Casadevall, A., and Dadachova, E. (2005) Radiological studies reveal radial differences in the architecture of the polysaccharide capsule of Cryptococcus neoformans. Eukaryotic Cell 4, 465–475

19. Gates, M. A., Thorkildson, P., and Kozel, T. R. (2004) Molecular architecture of the Cryptococcus neoformans capsule. Mol. Microbiol. 52, 13–24

20. Piccioni, M., Monari, C., Kenno, S., Pericolini, E., Gabrielli, E., Pietrella, D., Perito, S., Bistoni, F., Kozel, T. R., and Vecchiarelli, A. (2013) A purified capsular polysaccharide markedly inhibits inflammatory response during endotoxic shock. Infect. Immun. 81, 90–98

21. Makuuchi, K. (2010) Critical review of radiation processing of hydrogel and polysaccharide. Radiation Physics and Chemistry 79, 267–271

22. Jalan, N., Varshney, L., Misra, N., Paul, J., Mitra, D., Rairakhwada, D. D., Bhathena, Z., and Kumar, V. (2013) Studies on production of fructo-oligosaccharides (FOS) by gamma radiation processing of microbial levan. Carbohydr. Polym. 96, 365–370

23. Apar, D. K., and Özbek, B. (2008) Protein Releasing Kinetics of Bakers’ Yeast Cells by Ultrasound. undefined

24. Liu, D., Zeng, X.-A., Sun, D.-W., and Han, Z. (2013) Disruption and protein release by ultrasonication of yeast cells. Innovative Food Science & Emerging Technologies 18, 132–137

25. Joersbo, M., and Brunstedt, J. (1992) Sonication: A new method for gene transfer to plants. Physiol. Plant. 85, 230–234

26. Chemat, F., Zill-e-Huma, and Khan, M. K. (2011) Applications of ultrasound in food technology: Processing, preservation and extraction. Ultrason Sonochem 18, 813–835

27. Milner, H. W., Lawrence, N. S., and French, C. S. (1950) Colloidal dispersion of chloroplast material. Science 111, 633–634

28. Simpson, K. L., Wilson, A. W., Burton, E., Nakayama, T. O., and Chichester, C. O. (1963) Modified french press for the disruption of microorganisms. J. Bacteriol. 86, 1126–1127

29. Ait-Lahsen, H., Soler, A., Rey, M., de La Cruz, J., Monte, E., and Llobell, A. (2001) An antifungal exo-alpha-1,3-glucanase (AGN13.1) from the biocontrol fungus Trichoderma harzianum. Appl. Environ. Microbiol. 67, 5833–5839

30. Rosas, A. L., Nosanchuk, J. D., Gómez, B. L., Edens, W. A., Henson, J. M., and Casadevall, A. (2000) Isolation and serological analyses of fungal melanins. J. Immunol. Methods 244, 69–80

31. Dragotakes, Q., and Casadevall, A. (2018) Automated measurement of cryptococcal species polysaccharide capsule and cell body. J. Vis. Exp.

32. Wang, Y., Aisen, P., and Casadevall, A. (1996) Melanin, melanin “ghosts,” and melanin composition in Cryptococcus neoformans. Infect. Immun. 64, 2420–2424

33. Crawford, C. J., Cordero, R. J. B., Guazzelli, L., Wear, M. P., Bowen, A., Oscarson, S., and Casadevall, A. (2020) Exploring Cryptococcus neoformans capsule structure and assembly with a hydroxylamine-armed fluorescent probe. J. Biol. Chem. 295, 4327–4340

34. Zaragoza, O., Telzak, A., Bryan, R. A., Dadachova, E., and Casadevall, A. (2006) The polysaccharide capsule of the pathogenic fungus Cryptococcus neoformans enlarges by distal growth and is rearranged during budding. Mol. Microbiol. 59, 67–83

35. Cordero, R. J. B., Bergman, A., and Casadevall, A. (2013) Temporal behavior of capsule enlargement by Cryptococcus neoformans. Eukaryotic Cell 12, 1383–1388

36. Frases, S., Pontes, B., Nimrichter, L., Viana, N. B., Rodrigues, M. L., and Casadevall, A. (2009) Capsule of Cryptococcus neoformans grows by enlargement of polysaccharide molecules. Proc. Natl. Acad. Sci. USA 106, 1228–1233

37. Pierini, L. M., and Doering, T. L. (2001) Spatial and temporal sequence of capsule construction in Cryptococcus neoformans. Mol. Microbiol. 41, 105–115

38. Cordero, R. J. B., Pontes, B., Frases, S., Nakouzi, A. S., Nimrichter, L., Rodrigues, M. L., Viana, N. B., and Casadevall, A. (2013) Antibody binding to Cryptococcus neoformans impairs budding by altering capsular mechanical properties. J. Immunol. 190, 317–323

39. Maxson, M. E., Cook, E., Casadevall, A., and Zaragoza, O. (2007) The volume and hydration of the Cryptococcus neoformans polysaccharide capsule. Fungal Genet. Biol. 44, 180–186

40. Garcia-Rubio, R., de Oliveira, H. C., Rivera, J., and Trevijano-Contador, N. (2019) The fungal cell wall: candida, cryptococcus, and aspergillus species. Front. Microbiol. 10, 2993

41. Reese, A. J., and Doering, T. L. (2003) Cell wall alpha-1,3-glucan is required to anchor the Cryptococcus neoformans capsule. Mol. Microbiol. 50, 1401–1409

42. Reese, A. J., Yoneda, A., Breger, J. A., Beauvais, A., Liu, H., Griffith, C. L., Bose, I., Kim, M.-J., Skau, C., Yang, S., Sefko, J. A., Osumi, M., Latge, J.-P., Mylonakis, E., and Doering, T. L. (2007) Loss of cell wall alpha(1–3) glucan affects Cryptococcus neoformans from ultrastructure to virulence. Mol. Microbiol. 63, 1385–1398

43. Fonseca, F. L., Nimrichter, L., Cordero, R. J. B., Frases, S., Rodrigues, J., Goldman, D. L., Andruszkiewicz, R., Milewski, S., Travassos, L. R., Casadevall, A., and Rodrigues, M. L. (2009) Role for chitin and chitooligomers in the capsular architecture of Cryptococcus neoformans. Eukaryotic Cell 8, 1543–1553

44. Maxson, M. E., Dadachova, E., Casadevall, A., and Zaragoza, O. (2007) Radial mass density, charge, and epitope distribution in the Cryptococcus neoformans capsule. Eukaryotic Cell 6, 95–109

45. Cleare, W., and Casadevall, A. (1999) Scanning electron microscopy of encapsulated and non-encapsulated Cryptococcus neoformans and the effect of glucose on capsular polysaccharide release. Med Mycol 37, 235–243

46. Rakesh, V., Schweitzer, A. D., Zaragoza, O., Bryan, R., Wong, K., Datta, A., Casadevall, A., and Dadachova, E. (2008) Finite-element model of interaction between fungal polysaccharide and monoclonal antibody in the capsule of Cryptococcus neoformans. J. Phys. Chem. B 112, 8514–8522

47. Zaragoza, O., Taborda, C. P., and Casadevall, A. (2003) The efficacy of complement-mediated phagocytosis of Cryptococcus neoformans is dependent on the location of C3 in the polysaccharide capsule and involves both direct and indirect C3-mediated interactions. Eur. J. Immunol. 33, 1957–1967

48. Park, Y.-D., Shin, S., Panepinto, J., Ramos, J., Qiu, J., Frases, S., Albuquerque, P., Cordero, R. J. B., Zhang, N., Himmelreich, U., Beenhouwer, D., Bennett, J. E., Casadevall, A., and Williamson, P. R. (2014) A role for LHC1 in higher order structure and complement binding of the Cryptococcus neoformans capsule. PLoS Pathog. 10, e1004037

49. Zaragoza, O., and Casadevall, A. (2006) Monoclonal antibodies can affect complement deposition on the capsule of the pathogenic fungus Cryptococcus neoformans by both classical pathway activation and steric hindrance. Cell Microbiol. 8, 1862–1876

50. Clancy, C. J., Nguyen, M. H., Alandoerffer, R., Cheng, S., Iczkowski, K., Richardson, M., and Graybill, J. R. (2006) Cryptococcus neoformans var. grubii isolates recovered from persons with AIDS demonstrate a wide range of virulence during murine meningoencephalitis that correlates with the expression of certain virulence factors. Microbiology (Reading, Engl.) 152, 2247–2255

51. Dykstra, M. A., Friedman, L., and Murphy, J. W. (1977) Capsule size of Cryptococcus neoformans: control and relationship to virulence. Infect. Immun. 16, 129–135

52. Littman, M. L., and Tsubura, E. (1959) Effect of degree of encapsulation upon virulence of Cryptococcus neoformans. Proc Soc Exp Biol Med 101, 773–777

53. McFadden, D. C., De Jesus, M., and Casadevall, A. (2006) The physical properties of the capsular polysaccharides from Cryptococcus neoformans suggest features for capsule construction. J. Biol. Chem. 281, 1868–1875

54. BioRender [online] https://app.biorender.com/user/signin (Accessed November 12, 2021).

