## Supplemental Information for "*Cryptococcus neoformans* capsule regrowth experiments reveal dynamics of enlargement and architecture"

### Supporting Information:

Figure S1

Figure S2

Figure S3

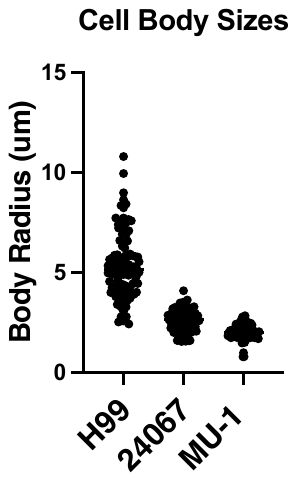

**Figure S1: Cell body size varies by *C. neoformans* strain.** Three strains were evaluated for cell body size using India ink images and the QCA assay. The cell bodies of single motif expressing strains 24067 and Mu-1 are smaller than those of mixed motif expressing strain H99.

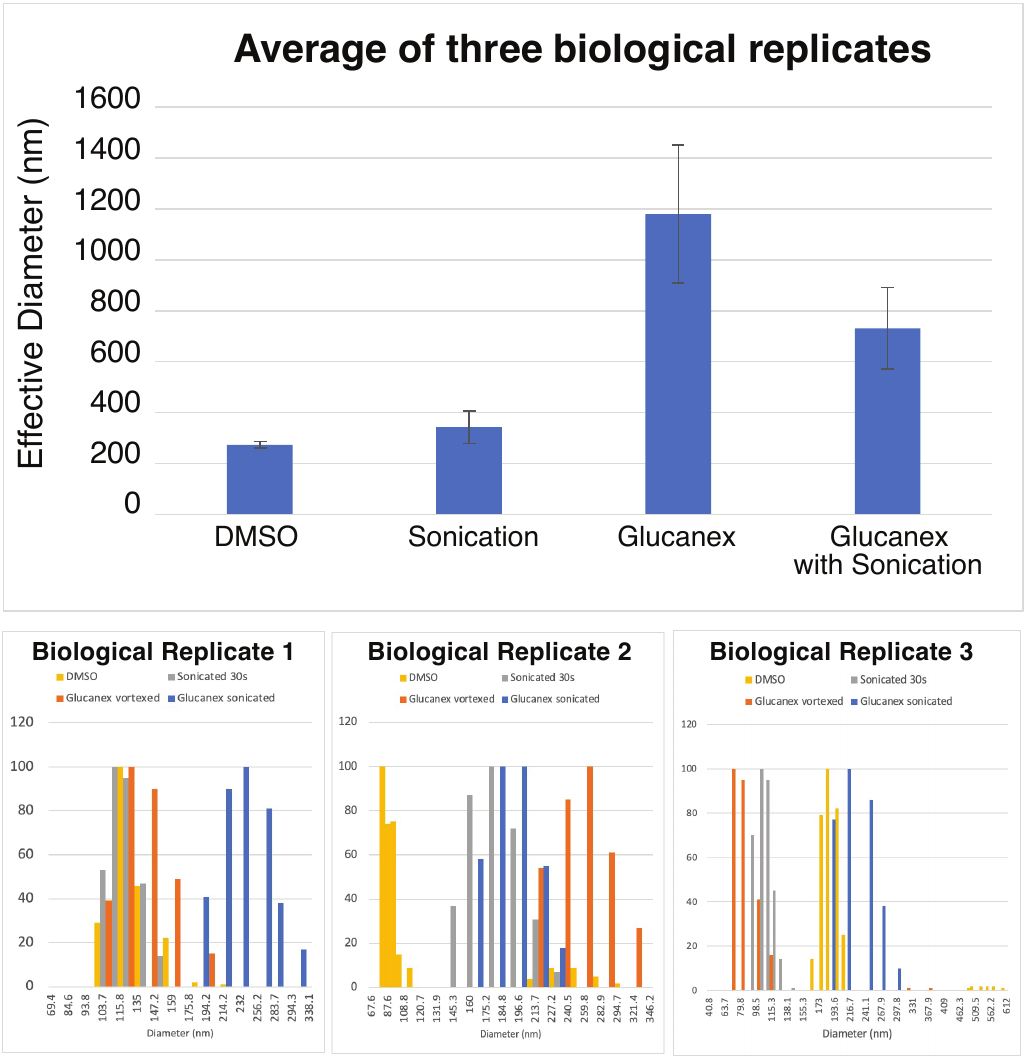

**Figure S2: DLS analysis of the size of particles released by CPS isolation methods. Top -** Average of three biological replicates for mean diameter shows a trend of larger particles from glucanex treatment than DMSO or sonication. **Bottom** – Full particle diameter range for each biological replicate shows high variability in size between both replicates and method of CPS isolation.

**Figure S3**: **Impact of lateral shear, and glucanex on capsular architecture A**. Immunofluorescence microscopy images of capsule ghosts produced by glucanex showing no cellular components (as stained by DAPI), but GXM as stained by 18B7-488. **B.** India ink stained Glucanex-derived capsule ghosts show blebbing of the capsule and loss of cell walls. **C.** Lateral shear generates capsule ghosts. Counterstained micrographs of encapsulated H99 *C. neformans* cells before (**a**) and after applying lateral shear pressure showing cell body and capsule dislocation (**b-c)** with scale bar representing 5 µm. Lateral sheared cells stained with Ubitex2B show India ink penetration of the capsule after treatment. Scale bar representing 100 µm. Cell body and capsule dislocation; capsule ghosts; broken cells; cell aggregates; capsule fragments.  All images adjusted for brightness and contrast in the same manner.
